# An efficient single-cell transcriptomics workflow to assess protist diversity and lifestyle

**DOI:** 10.1101/782235

**Authors:** Henning Onsbring, Alexander K. Tice, Brandon T. Barton, Matthew W. Brown, Thijs J. G. Ettema

## Abstract

Most diversity in the eukaryotic tree of life are represented by microbial eukaryotes, which is a polyphyletic group also referred to as protists. Among the protists, currently sequenced genomes and transcriptomes give a biased view of the actual diversity. This biased view is partly caused by the scientific community, which has prioritized certain microbes of biomedical and agricultural importance. Additionally, it is challenging to establish protist cultures, which further influence what has been studied. It is now possible to bypass the time-consuming process of cultivation and directly analyze the gene content of single protist cells. Single-cell genomics was used in the first experiments where individual protists cells were genomically explored. Unfortunately, single-cell genomics for protists are often associated with low genome recovery and the assembly process can be complicated because of repetitive intergenic regions. Sequencing repetitive sequences can be avoided if single-cell transcriptomics is used, which only targets the part of the genome that is transcribed. In this study we test different modifications of Smart-seq2, a single-cell RNA sequencing protocol originally developed for mammalian cells, to establish a robust and more cost-efficient workflow for protists. The diplomonad *Giardia intestinalis* was used in all experiments and the available genome for this species allowed us to benchmark our results. We could observe increased transcript recovery when freeze-thaw cycles were added as an extra step to the Smart-seq2 protocol. Further we tried reducing the reaction volume and purifying with alternative beads to test different cost-reducing changes of Smart-seq2. Neither improved the procedure, and cutting the volumes by half actually led to significantly fewer genes detected. We also added a 5’ biotin modification to our primers and reduced the concentration of oligo-dT, to potentially reduce generation of artifacts. Except adding freeze-thaw cycles and reducing the volume, no other modifications lead to a significant change in gene detection. Therefore, we suggest adding freeze-thaw cycles to Smart-seq2 when working with protists and further consider our other modification described to improve cost and time-efficiency.

## Background

Protists are undersampled among the eukaryotes in terms of genome and transcriptome sequencing efforts, the scientific community has mainly generated such data for plants, fungi, and animals [1]. Generation of genome and transcriptome data for protists is challenging, since only a small minority of this group have been cultivated under controlled laboratory conditions [2-4]. Methods that are using only a single cell as input can bypass the time-consuming work of establishing a culture. Single-cell genomics is an example of such an approach, which has been applied to expand our knowledge about protist diversity. However, attempts to sequence the genome from single protist cells are often associated with poor genome recovery [5-7]. Another possibility to generate gene content data from uncultivated protists is single-cell RNA sequencing, which never targets the repetitive intergenic regions that can cause trouble with genomic approaches.

Single-cell RNA sequencing was first tested on protists in a study from 2014 [8] that used the commercial SMARTer kit, achieving a result comparable to conventional sequencing based on RNA extraction from a culture. However, the cells ranged from 50 to 500 µm in size that were analyzed in this study. Single-cell RNA sequencing of a haptophyte and dinoflagellate (8 and 15 µm cell size respectively) were later tested in 2017 by Liu et al. [9], where an updated version of the SMARTer kit (SMART-Seq) was used. In this study only 3% of the transcripts were recovered on average for the haptophyte and 15% for the dinoflagellate. Modifications of the SMART-Seq protocol might be needed to achieve better results for smaller cells. Unfortunately, modifications of the procedure can be complicated when a commercial kit is used, especially since some of the components tend to be kept undisclosed and the kits themselves are expensive per reaction.

In this study we have instead used Smart-seq2 [10] as a starting point, which is fully based on off-the-shelf reagents and performs better than the SMARTer kit, both when it comes to gene detection and coverage [11]. Unlike Liu et al., we have not performed any RNA extraction prior to cDNA synthesis, which could potentially reduce transcript recovery. In the Smart-seq2 workflow we have tested different changes, which might improve the generation of cDNA from smaller protists. Our modifications of Smart-seq2 offer improved lysis and less dependence on quality control compared to the original protocol. We have benchmarked all protocols tested in this study on *Giardia intestinalis*, which genome is sequenced [12]. The problem that mainly has to be addressed when working with environmental protists is lysis of the cell. Also for cells with low RNA content, there can be a problem with unspecific amplification due to changed balance between the concentration of oligos and mRNA of the cell [13]. The potential problem with lysis is addressed by using freeze-thaw cycles in −80 °C chilled isopropanol, which previously have been reported as a successful lysis procedure [14, 15]. Besides the improved lysis we already know can be crucial, we test modifications of Smart-seq2 that could make the protocol more cost-efficient or reduce the generation of artifacts during cDNA synthesis.

## Results

### Gene Detection and Coverage

Single *G. intestinalis* trophozoites were sorted using fluorescence-activated cell sorting and seven different protocols for generation of transcriptomes were applied, including Smart-seq2 and modified versions of Smart-seq2 (Fig. 1). Freeze-thaw cycles were added to all six modifications of Smart-seq2. Additionally, five of the modified versions of Smart-seq2 had one or all of the following changes: biotinylated 5’ end of primers, other beads for cDNA purification, lower reaction volume and less oligo-dT primers than Smart-seq2 (see methods for details).

**Fig. 1.**
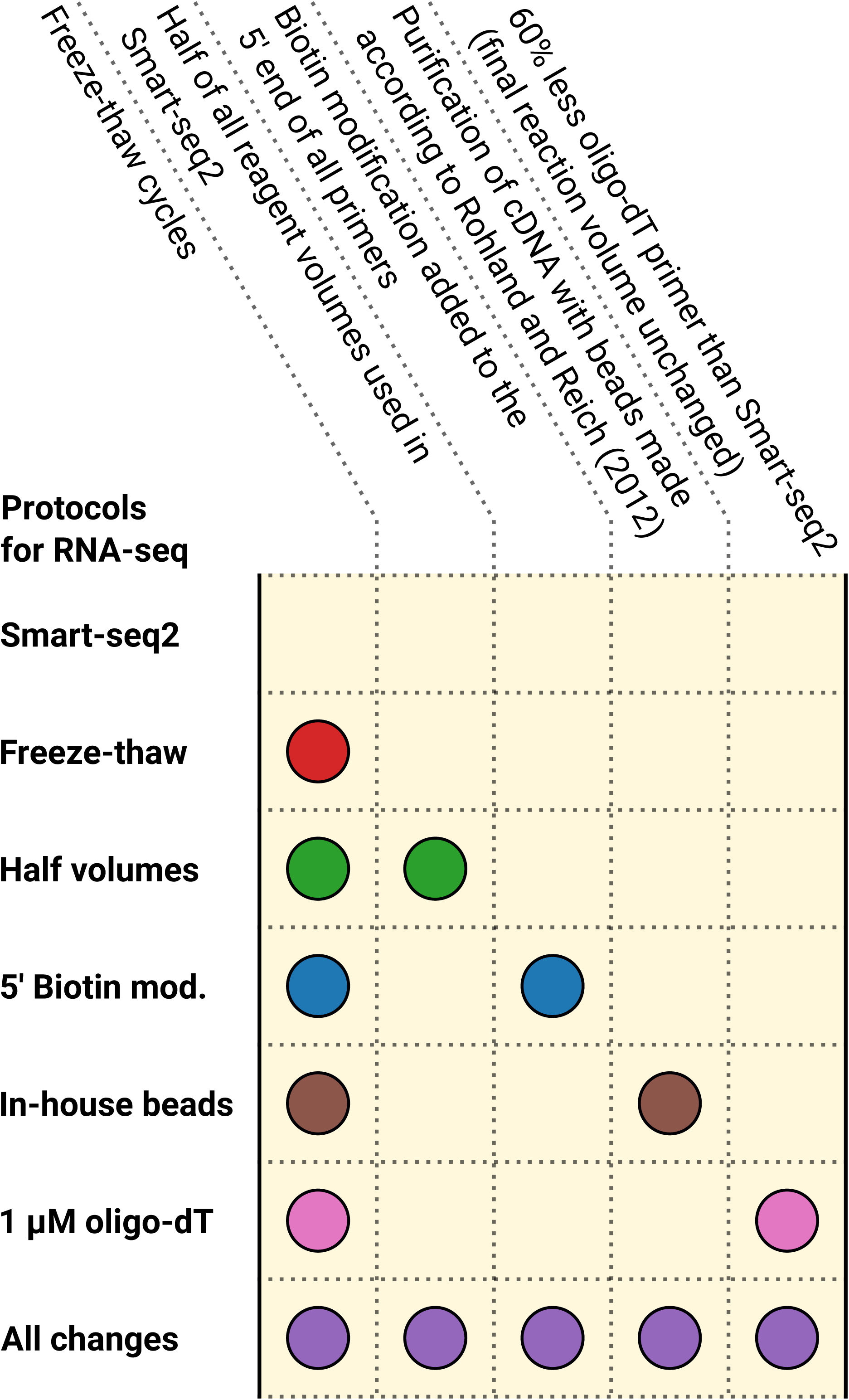
Overview of how the protocols tested in this study differ from Smart-seq2.

The sequencing data generated from all transcriptomes corresponded to 703 Gb, covering 55 individual cells. We detected on average 4524 to 4992 genes in all tested protocols (Fig. 2a), representing 70-77% of the total protein coding genes in the genome of *G. intestinalis* [12]. Using fragments per kilobase of transcript per million mapped reads (FPKM) allowed us to take the abundance of transcripts into consideration in our analysis. All protocols, except the version where all tested changes are implemented, differ by only one treatment compared to Smart-seq2 with freeze-thaw cycles. Therefore, we used this “Freeze-thaw” protocol as the point of reference in our pairwise comparisons. Using the unmodified Smart-seq2 lead to significantly fewer genes being detected among the medium and high abundance transcripts (FPKM > 0.1 and > 1). When half volumes of the standard reagents were used throughout the protocol, significantly fewer genes were detected for both low and medium abundance transcripts (FPKM > 0 and > 0.1).

**Fig. 2.**
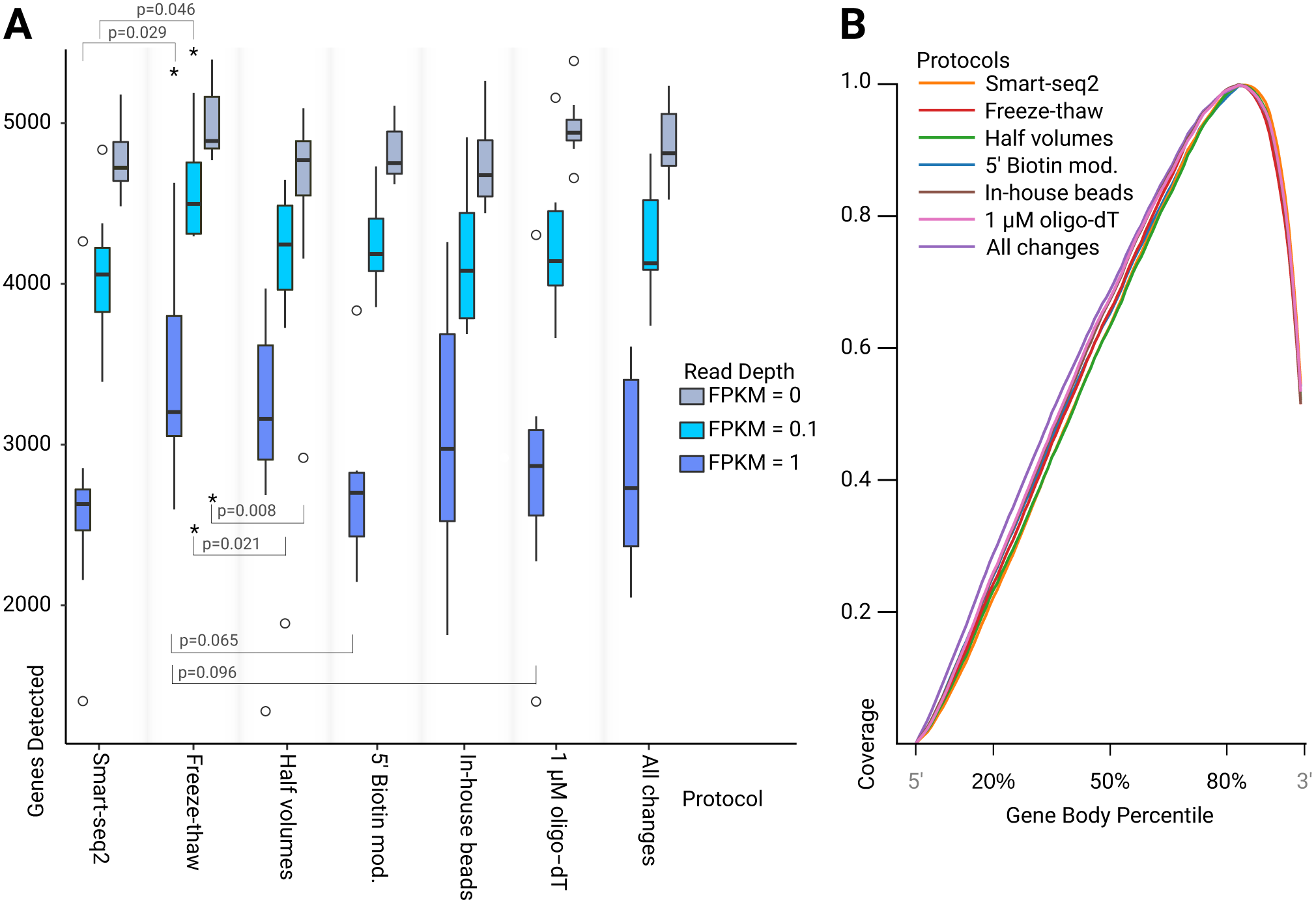
Transcriptome quality statistics. **a** Box and whisker plot showing number of genes detected for all permutations at three expression levels. An asterix indicates significance when compared to our “freeze-thaw” protocol (p < 0.05). P-values below plots indicate that the performance of the treatment was worse at gene detection. **b** Average gene body coverage for all genes detected in the *G. intestinalis* genome by each protocol.

We also tested the use of biotinylated primers, reduced concentration of oligo-dT primers, beads made in-house or a combination of all modifications of Smart-seq2 tested in this study, neither of these protocols performed significantly different from the “Freeze-thaw” protocol. However, we saw a marginal decreases in gene detection when using 1 µM oligo-dT (glm, p = 0.096), and biotinylated primers (glm, p = 0.065) at a read depth of FPKM > 1 (see Table 1). Unmodified Smart-seq2, as well as all our modified protocols, show a 3’ bias in gene body coverage (Fig. 2b). This bias is common to protocols that use oligo-dT priming during cDNA synthesis [16].

**Table 1.**
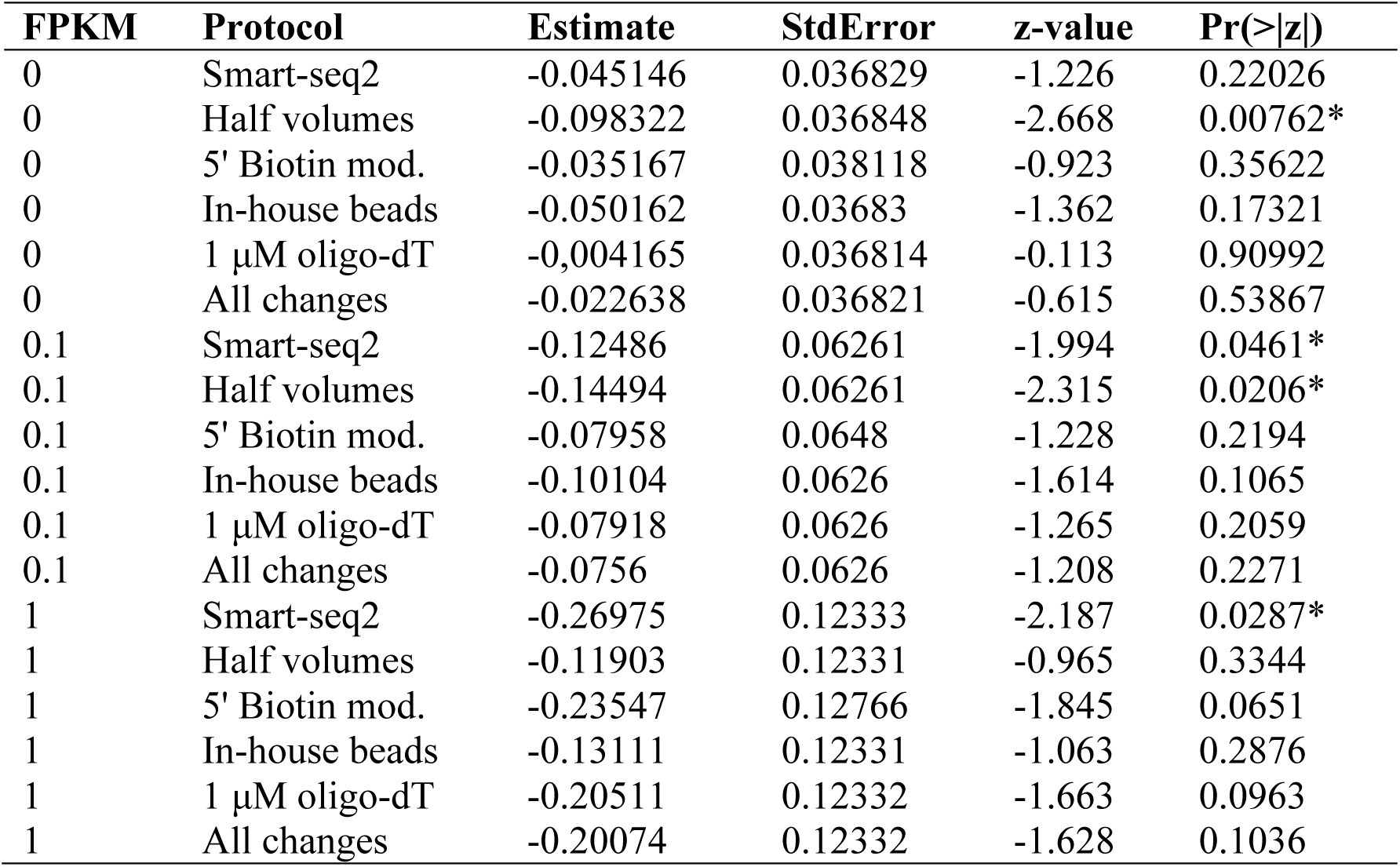
Comparison of the number of genes detected for Smart-seq2, and five modified Smart-seq2 variants, against our “Freeze-thaw” protocol using a generalized linear model with a negative binomial error distribution. Significant p-values are indicated with an asterisk.

| **FPKM** | **Protocol** | **Estimate** | **StdError** | **z-value** | **Pr(>\|z\|)** |
| --- | --- | --- | --- | --- | --- |
| 0 | Smart-seq2 | -0.045146 | 0.036829 | -1.226 | 0.22026 |
| 0 | Half volumes | -0.098322 | 0.036848 | -2.668 | 0.00762* |
| 0 | 5' Biotin mod. | -0.035167 | 0.038118 | -0.923 | 0.35622 |
| 0 | In-house beads | -0.050162 | 0.03683 | -1.362 | 0.17321 |
| 0 | 1 μM oligo-dT | -0.004165 | 0.036814 | -0.113 | 0.90992 |
| 0 | All changes | -0.022638 | 0.036821 | -0.615 | 0.53867 |
| 0.1 | Smart-seq2 | -0.12486 | 0.06261 | -1.994 | 0.0461* |
| 0.1 | Half volumes | -0.14494 | 0.06261 | -2.315 | 0.0206* |
| 0.1 | 5' Biotin mod. | -0.07958 | 0.0648 | -1.228 | 0.2194 |
| 0.1 | In-house beads | -0.10104 | 0.0626 | -1.614 | 0.1065 |
| 0.1 | 1 μM oligo-dT | -0.07918 | 0.0626 | -1.265 | 0.2059 |
| 0.1 | All changes | -0.0756 | 0.0626 | -1.208 | 0.2271 |
| 1 | Smart-seq2 | -0.26975 | 0.12333 | -2.187 | 0.0287* |
| 1 | Half volumes | -0.11903 | 0.12331 | -0.965 | 0.3344 |
| 1 | 5' Biotin mod. | -0.23547 | 0.12766 | -1.845 | 0.0651 |
| 1 | In-house beads | -0.13111 | 0.12331 | -1.063 | 0.2876 |
| 1 | 1 μM oligo-dT | -0.20511 | 0.12332 | -1.663 | 0.0963 |
| 1 | All changes | -0.20074 | 0.12332 | -1.628 | 0.1036 |

### Identification of phylogenetic markers

To obtain a rough estimate how much data is needed to be able to extract marker genes to build a multi-gene concatenated alignment for phylogenomics of an unknown protist, we down-sampled our data and ran multiple *de novo* assemblies in several iterations (Fig. 3). Generally among the comparisons, based on different number of reads used in the assembly, we observe that Smart-seq2 with freeze-thaw cycles identified significantly more markers than Smart-seq2, 1 µM oligo-dT, 5’ biotin modification and when all changes where applied. As a proxy for a phylogenomic analysis dataset, we calculated the number of observed BUSCO in the Eukaryota odb9 dataset. The number of BUSCO markers detected did not increase much if more sequencing data was generated beyond 500 thousand read pairs, which correspond to 150 Mbp sequencing data. This indicates that a low amount of data is needed if the only goal is to find markers for a phylogenomic analysis.

**Fig. 3.**
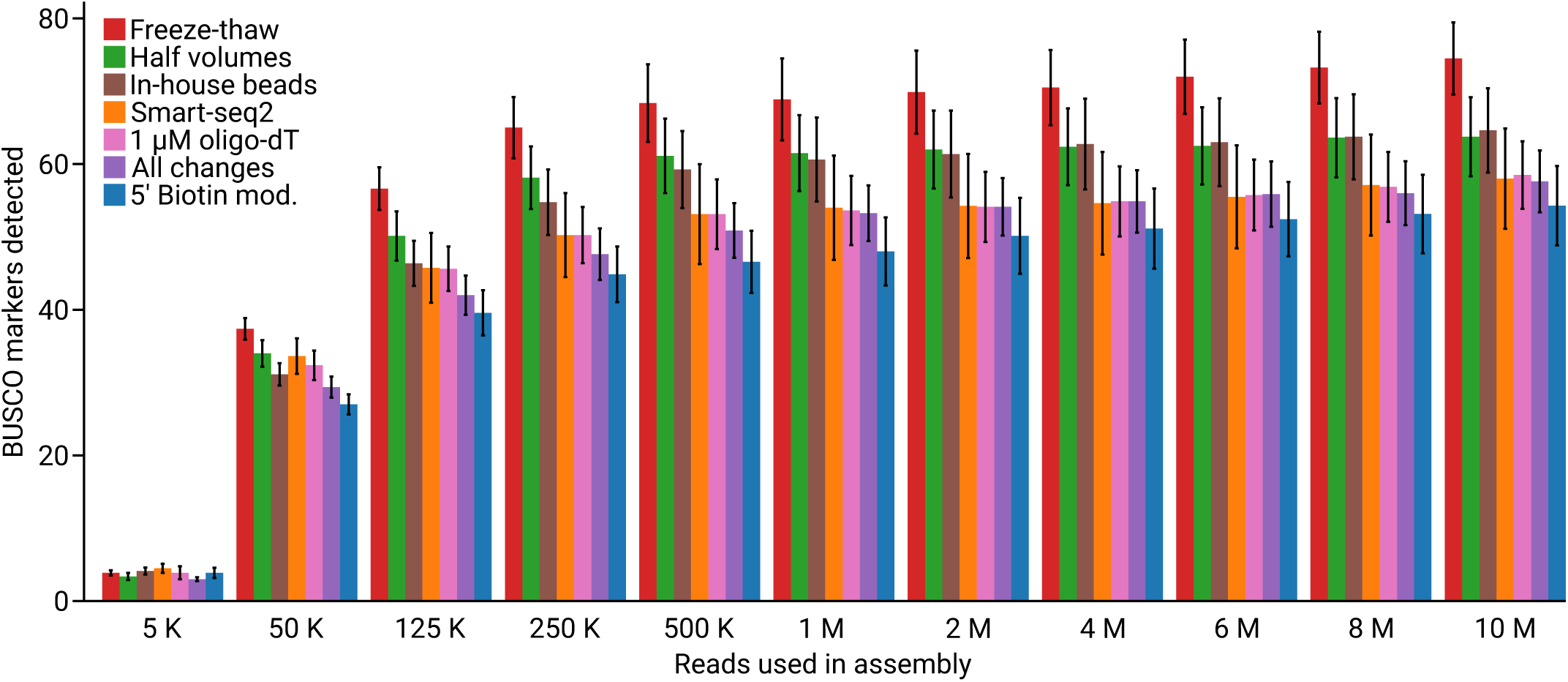
Number of BUSCO markers found in *de novo* assemblies based on different amount of data. Bars represent the average number of BUSCO markers found in the *de novo* assemblies, error bars represent standard error. The average number of BUSCO markers found varied between 54 to 75 (out of the 303 proteins in the eukaryota_odb9 dataset). Retrieving around 54 to 75 markers from our single-cell transcriptomes can be compared to the reference transcriptome of *G. intestinalis* on NCBI, which encode 146 of the BUSCO markers from eukaryota_odb9.

## Discussion

By performing Smart-seq2, and six alternative modifications of this protocol, we generated 55 transcriptomes of single *G. intestinalis* cells. The raw sequencing reads allowed us to generate statistics for gene detection and gene body coverage by mapping to the *G. intestinalis* genome [12]. Our experiment shows that adding six freeze-thaw cycles to the Smart-seq2 protocol will not decrease the RNA quality in a way that negatively affect gene detection or gene body coverage. Adding these freeze-thaw cycles actually turned out to significantly increase the number of genes detected among the two highest read depths analyzed. Because of this improvement and since we expect that many protists are harder to lyse than the mammalian cells used to optimized Smart-seq2, we suggest that freeze-thaw cycles should be used when generating protist transcriptomes from single-cell input. Because our experience is that the freeze-thaw cycles can be necessary to get a successful cDNA library [14], we have used freeze-thaw in all of our modifications of the Smart-seq2 protocol. Therefore, Smart-seq2 with freeze-thaw cycles becomes the point of reference and will be used as our control in pairwise comparisons to other tested protocols.

The only modified version of Smart-seq2 we tested in this study, that lead to significantly fewer genes detected, was when we reduced all reagent volumes to half of what is used in the original protocol. The lower performance could be due to the unfavorable change in ratio between reaction volume and surface area of the test tube wall, which can absorb nucleic acid [17]. Despite the lower performance, cutting all volumes by half may be considered in experimental design due to cost savings associated with using less reagents, which could be important when running many reactions.

It has been reported that modifications of Smart-seq2 is necessary when working with cells with extremely low RNA content, e.g. concatamerization of the template switching oligo can prevent the generation of usable cDNA libraries (Picelli 2016). To prevent such generation of background during cDNA synthesis we tried adding a 5’ biotin modification for all primers, which is also recommended in an updated version of the Smart-seq2 [18]. Adding the 5’ biotin modification did not increase the number of genes detected and the number of BUSCO markers were significantly fewer than what was recovered from the control. At the same time when the biotin modification was not used, the potential problem with concatamers [19] was never observed (Fig. S1). Based on recommendations from other studies this option can be considered as an insurance against failed cDNA generation, especially for cells with lower RNA content than *G. intestinalis*.

Another protocol modification that could reduce the amount of artifacts is changing the concentration of primers, which we tested by decreasing the concentration of oligo-dT by 60%. This was done since the imbalance of primers and mRNA has been claimed to be one of the reasons why background is generated when working with cells that have low mRNA content [13]. Reducing the concentration of oligo-dT with 60% did not increase the number of genes detected, and significantly less BUSCO markers were found. Therefore, using less oligo-dT should not be considered for cells with as much RNA as *G. intestinalis* or more, if the goal is to maximize transcript recovery.

Besides the previously discussed oligo-concatamers, an artifact that we did see in our fragment length analysis was the formation of primer dimers. We could reduce the amount of primer dimers by preparing beads for purification of the amplified cDNA (Fig. 4). However, we did not observe any aspect of the protocol that improved by this change, except lower cost of consumables for DNA purification compared to Smart-seq2.

**Fig. 4.**
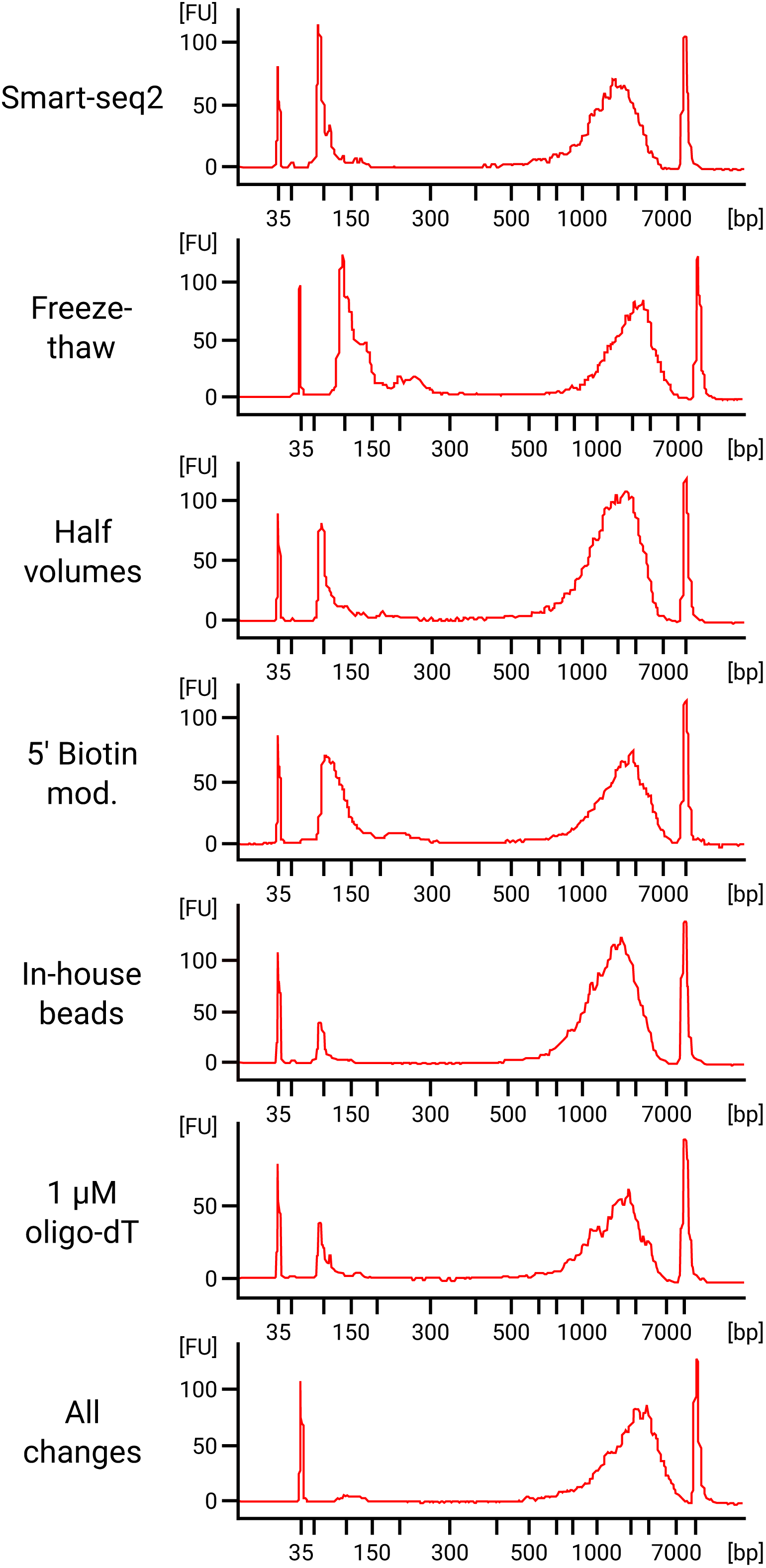
Fragment length analysis of seven cDNA libraries covering each variation of the single-cell RNA sequencing protocol tested.

If a high number of transcriptomes are going to be generated, we recommend using all modifications of Smart-seq2 tested in this study. However, the “All changes” protocol did not lead to higher transcript recovery compared to the control. The important benefit of the “All changes” workflow is that the user becomes less dependent on the time-consuming and costly fragment length analysis step. When generating many transcriptomes it is advantageous to be able to identify failed reactions by just measuring the DNA concentration. If all modifications tested in this study are applied all at once, then the failed reactions will typically measure well below the lowest recommended input for sequencing library preparation. Therefore this will save time and money by reducing the need for fragment length analysis, while also less reagents and cheaper purification beads are used. However, checking the fragment length distribution on a subset of the generated cDNA libraries is always recommended. Fragment length analysis allows detection of ribonuclease contamination and can prevent the user from proceeding to the next step in the workflow with a degraded sample. If there is no equipment available for detailed fragment length analysis, or if the user wants to reduce cost, additional amplification of the sequencing libraries combined with gel electrophoresis has previously been used as an alternative [20].

## Conclusions

Our results from testing seven different protocols for generation of cDNA suggests that freeze-thaw cycles should be added to a single-cell transcriptomics workflow for protists. To save money, all volumes in Smart-seq2 can be reduced to half and lab-prepared purification beads can be used, but neither of these changes leads to any improvements in gene detection. Actually, using half of the recommended Smart-seq2 volumes might reduce the transcript recovery. A 5’ biotin modification of the primers can be considered as an insurance against concatamers, but this change could be at the expense of lower transcript recovery as well.

To become less dependent on quality control, all changes tested in this study can simultaneously be applied in one protocol. The dependency on quality control is reduced since failed reactions will have a cDNA concentration close to 0, and therefore it is possible to discard unsuccessful cDNA libraries only based on DNA concentration.

Transcriptomes encoding markers for multi-gene concatenated phylogenies can be generated with single-cell RNA sequencing, even with low amount of sequencing data. All variations of Smart-seq2 tested in this study are suitable options for generation of data to perform phylogenomic analysis. Therefore, instead of optimizing transcript recovery, factors such as time or cost-efficiency can be considered.

## Methods

### Cell sorting

Trophozoites of *Giardia intestinalis*, isolate WB, were grown to confluence in 10 mL flat bottom tubes (NUNC) and detached on ice for 10 minutes. The cell suspension was transferred to a 15 mL Falcon tube and centrifuged at 500 *g* for 10 min. The supernatant was discarded and resuspended in 500 µL 1xPBS. Prior to sorting, the sample was prepared using a cell suspension of harvested trophozoites diluted 10 times in sterile filtered 1xPBS and stained with DAPI and Propidium Iodide (PI) to a final concentration of 1 µg/mL and 200 nM respectively for 10 min. The sorting was performed with a MoFlo Astrios EQ (Beckman Coulter, USA) flow cytometer using the 355 and 532 nm lasers for excitation, a 100 µm nozzle, sheath pressure of 25 psi and 0.2 µm filtered 1xPBS as sheath fluid. Live cells were identified using scatter properties in combination with a singlets gate and exclusion of dead PI positive cells. Individual cells were deposited into 12 × 8-well strips containing 2.3 µl or 4.3 µl of lysis buffer using a CyCloneTM robotic arm and the most stringent single cell sort settings (e.g single mode, 0.5 drop envelope).

The lysis buffer were for some reactions prepared according to Smart-seq2 [10], and altered in some of the modified versions of the protocol (see the methods paragraph “cDNA synthesis” for details). A UV-laser (355 nm) was used for excitation of DAPI and emission was collected by a 448/59 nm filter. Excitation of PI and collection of emitted light was done with a 532 nm laser with a 622/22 nm filter. Side scatter was used as trigger channel. The plate and sample holder were kept at 4 °C during the sort. The 8-strips were sorted two by two, quickly spun down and temporarily stored at −20 °C until the sort was finished before transfer to a −80 °C freezer.

### cDNA synthesis

The cDNA was prepared according to Smart-seq2 [10], and six modified versions of Smart-seq2, using 24 cycles of cDNA amplification in each case. We generated 8 cDNA libraries for every version of the protocol. All six modified versions of Smart-seq2 included freeze-thaw cycles as an extra lysis step. The freeze-thaw cycles were performed by first thawing the frozen cells in room-tempered water for 10 s directly after taken out of the freezer. Immediately after the 10 s thaw, the tubes were frozen down again in −80 °C isopropanol for 10 s. This freeze-thaw cycle was repeated six times.

The specific changes applied for each of the six protocols were 1) No additional changes to Smart-seq2 besides the freeze-thaw cycles. 2) Decreasing the oligo-dT primer concentration to 1 µM, instead of 2.5 µM, in the first mix of primer, dNTP and lysis that is added to the cell. 3) Using the beads described by N Rohland and D Reich [21], with a 17% PEG concentration, for purification of the amplified cDNA. 4) All volumes were reduced to half of what is used in the original Smart-seq2 protocol. 5) Adding a 5’ biotin modification to all primers, including the one used for template switching. 6) Using all these changes at once, including freeze-thaw cycles, decreased oligo-dT concentration, using the beads made in-house, cutting all volumes by half and 5’ biotin modification added to primers.

### Tagmentation and sequencing

DNA concentration was measured with Qubit dsDNA HS Assay Kit (Thermo Fisher Scientific). Fragment length analysis was done using Agilent High Sensitivity DNA Kit with a 2100 Bioanalyzer Instrument on a subset of the purified cDNA (Fig. S1). The purified cDNA was then diluted so each sequencing library preparation reaction had a 1.3 ng input of DNA, followed by using the Nextera XT DNA Library Preparation Kit (Illumina). One Nextera XT library failed, leading to that only 7 replicates based on the protocol using 5’ biotinylated primers were included in the sequencing run. A total of 55 single-cell transcriptomes were sequenced on a separate lane of Illumina NovaSeq S4 (2 × 150 bp reads).

### Read Mapping and Quantification

Sequencing data quality was assessed using FastQC v0.11.8 [22] and visualized using MultiQC [23]. Low quality bases and adaptors were removed using Trimmomatic v0.39 with the options “ILLUMINACLIP: 2:30:10 LEADING:5 TRAILING:5 SLIDINGWINDOW:5:16 MINLEN:60” [24]. Reads were then mapped to the *G. intestinalis* genome (GCF_000002435.1) using tophat2 with default settings [25]. The python scripts geneBody_coverage.py and FPKM_count.py from RSeQC-2.6.4 were used to examine read distribution across genes and calculate FPKM values for all libraries respectively [26]. The box and whisker plot for number of genes detected was generated in R using the ggplot package. While the line graph showing gene body coverage was made using matplotlib via a custom python script.

### Statistical Analyses

We compared the number of genes detected at three expression/abundance levels (FPKM > 0, > 0.1, >1) for unmodified Smart-seq2 and five protocol variants against our “Freeze-thaw” protocol. We used a generalized linear model with a negative binomial error distribution to correct for overdispersion. All statistical analyses were conducted with the *glm* module in R.

### BUSCO analysis

Separate assemblies were done for each cell using Trinity v2.4.0 [27]. For every cell we assembled 11 different assemblies using the following number of reads as input: 10 million, 8 million, 6 million, 4 million, 2 million, 1 million, 500 thousand, 250 thousand, 125 thousand, 50 thousand, 5 thousand. Each assembly was then analyzed with BUSCO v3.1.0 [28], using the eukaryota_odb9 dataset. A BUSCO analysis was also done on the reference transcriptome of *G. intestinalis* from the NCBI database.

## List of abbreviations

BUSCO: Benchmarking universal single-copy orthologs
FPKM: Fragments per kilobase of transcript per million mapped reads

## Availability of data and material

All raw sequencing reads generated in this study have been submitted to NCBI under the BioProject PRJNA545787.

## Authors’ contributions

H.O. and T.J.G.E. conceived and planned the study. H.O. generated the cDNA. H.O., A.K.T. and B.T.B. analyzed the data. H.O., A.K.T., M.W.B. and T.J.G.E. interpreted the results and wrote the manuscript. All authors read, edited, and approved the final manuscript.

## Acknowledgements

We want to thank Showgy Ma’ayeh (Uppsala University) for providing *G. intestinalis* cells. Single *G. intestinalis* cells were sorted by the Microbial Single Cell Genomics Facility (SiCell), which is a part of Science for Life Laboratory at Uppsala University. All sequencing was performed by the National Genomics Infrastructure sequencing platforms at the Science for Life Laboratory at Uppsala University, a national infrastructure supported by the Swedish Research Council (VR-RFI) and the Knut and Alice Wallenberg Foundation.

## Figures, tables and additional files

**Fig. S1** Fragment length analysis of cDNA libraries covering multiple replicates from each variant of the protocol tested.

