## Supplementary figures and images for "An efficient single-cell transcriptomics workflow to assess protist diversity and lifestyle"

### Supplemental Figure 1

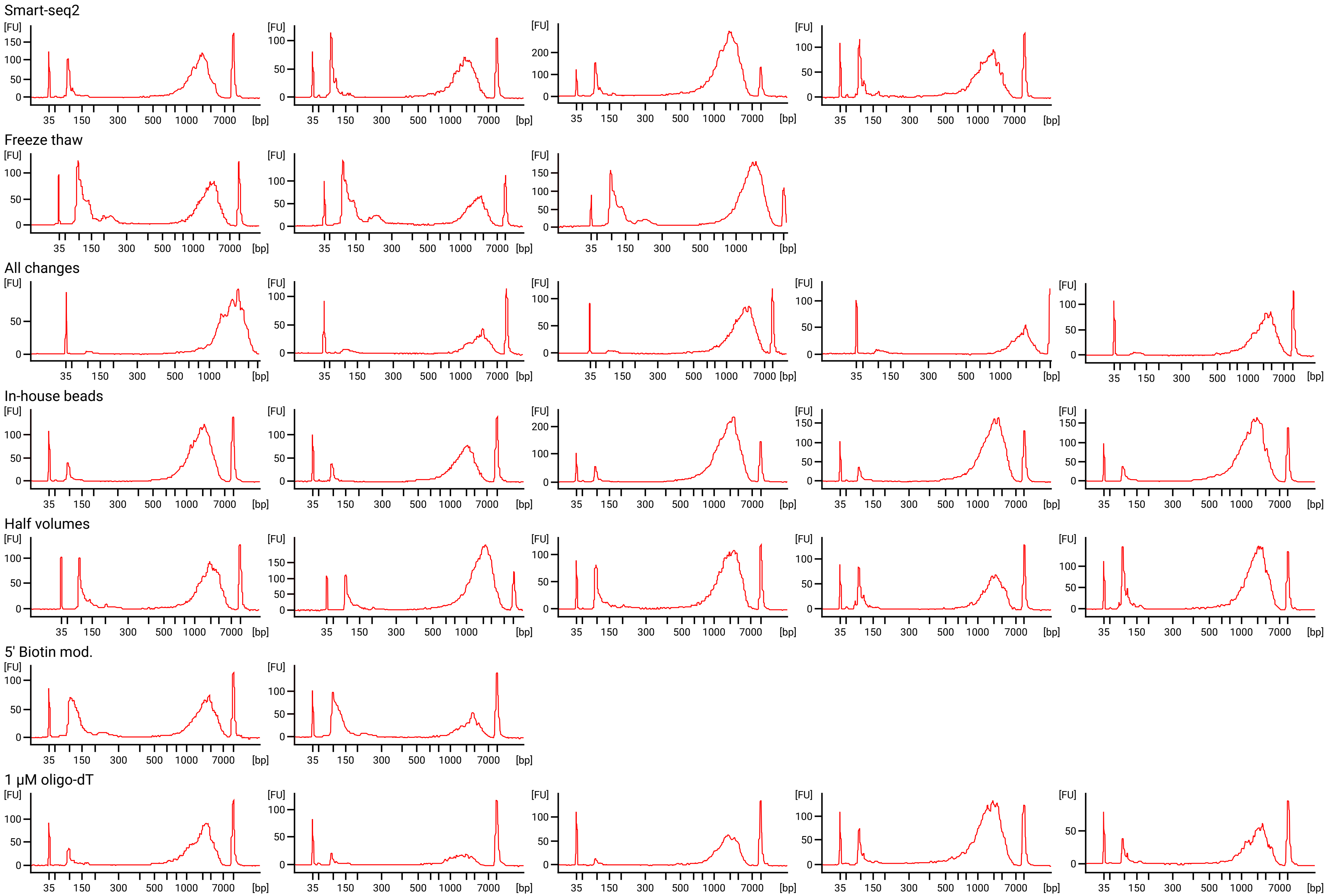
